# Spatial Coherence in DNA Barcode Networks

**DOI:** 10.1101/2024.05.12.593725

**Authors:** David Fernandez Bonet, Johanna I. Blumenthal, Shuai Lang, Simon K Dahlberg, Ian T. Hoffecker

## Abstract

Sequencing-based microscopy is a novel, optics-free method for imaging molecules in biological samples using molecular DNA barcodes, spatial networks, and sequencing technologies. Despite its promise, the principles determining how these networks preserve spatial information are not fully understood. Current validation methods, which rely on comparing reconstructed positions to expected results, would benefit from a deeper understanding of these principles. Here, we introduce the concept of spatial coherence— a set of fundamental properties of spatial networks that quantifies the alignment between topological relationships and Euclidean geometry. Our findings show that spatial coherence is an effective method for evaluating a network’s capacity to maintain spatial fidelity and identify distortions, independent of prior information. This framework provides a cost-effective validation tool for sequencing-based microscopy by taking advantage of the fundamental properties of spatial networks in nanoscale systems.

---

Spatial molecular mapping methods in tissues are expanding our understanding of molecular physiology, development, and pathology Lewis et al. (2021); Moses and Pachter (2022). These include optical molecular imaging methods, offering finely resolved images for multiple simultaneous targets, whereas single-cell methods (Shapiro et al., 2013; Gawad et al., 2016) provide data for a large number of genes but lack spatial information about where cells are in the tissue. Spatial omics techniques (Ke et al., 2013; Lee et al., 2015; Ståhl et al., 2016; Wang et al., 2018; Karaiskos et al., 2017; Satija et al., 2015; Achim et al., 2015; Halpern et al., 2017; Rodriques et al., 2019) seek to eliminate this trade-off with high multiplicity coupled with spatial resolution, but are also constrained by instrumentation-demanding bottleneck steps for generating spatial reference maps (e.g. by microscopy-based *in situ* sequencing or array printing) to look up molecule locations.

Sequencing-based microscopy(Glaser et al., 2015; Schaus et al., 2017; Boulgakov et al., 2018; Weinstein et al., 2019; Hoffecker et al., 2019; Gopalkrishnan et al., 2020; Boulgakov et al., 2020; Greenstreet et al., 2022; Karlsson et al., 2024), in contrast, is an emerging family of techniques that seeks to spatially resolve many molecular targets but without a spatial reference map, relying instead only on DNA to capture spatial information in the structure of sequencing data itself. This “inside-out” approach instead of array-based methods offers potential advantages, including molecular target multiplexing, scalability, reduced instrumentation, and isotropic imaging in 3D or optically opaque samples.

Sequencing-based microscopy methods (Fig.1a) exploit network capacity to capture spatial information. Initially, molecules of interest –such as messenger RNA or proteins from a biological sample– are tagged with unique DNA identifiers (UMIs or barcodes). These barcodes are clonally amplified to form polonies, or local patches of uniquely identifiable DNA. Each polony interacts with neighboring polonies via covalent linkage, forming a proximity graph (Mathieson and Moscato, 2019) (here-after called a spatial network). Spatial networks encapsulate individual molecular identities as nodes and spatial information in the form of network structure. After isolating and sequencing the DNA from the sample, this information can be extracted and used to reconstruct the network and thereby the molecule spatial positions. Different sequencing-based microscopy modalities have been modeled *in silico* (Hoffecker et al., 2019; Bonet and Hoffecker, 2023; Greenstreet et al., 2022; Glaser et al., 2015; Boulgakov et al., 2020) and implemented experimentally (Weinstein et al., 2019; Karlsson et al., 2024; Qian and Weinstein, 2023). Although each modality comes with a model of how networks form, e.g. saturated polony forests that form tessellations or beacon-receiver communication via diffusion, these models are ultimately idealized representations of the network formation chemistry. Real data can contain spurious interactions that do not correlate with spatial relationships, impacting reconstruction potential.

**Figure 1.**
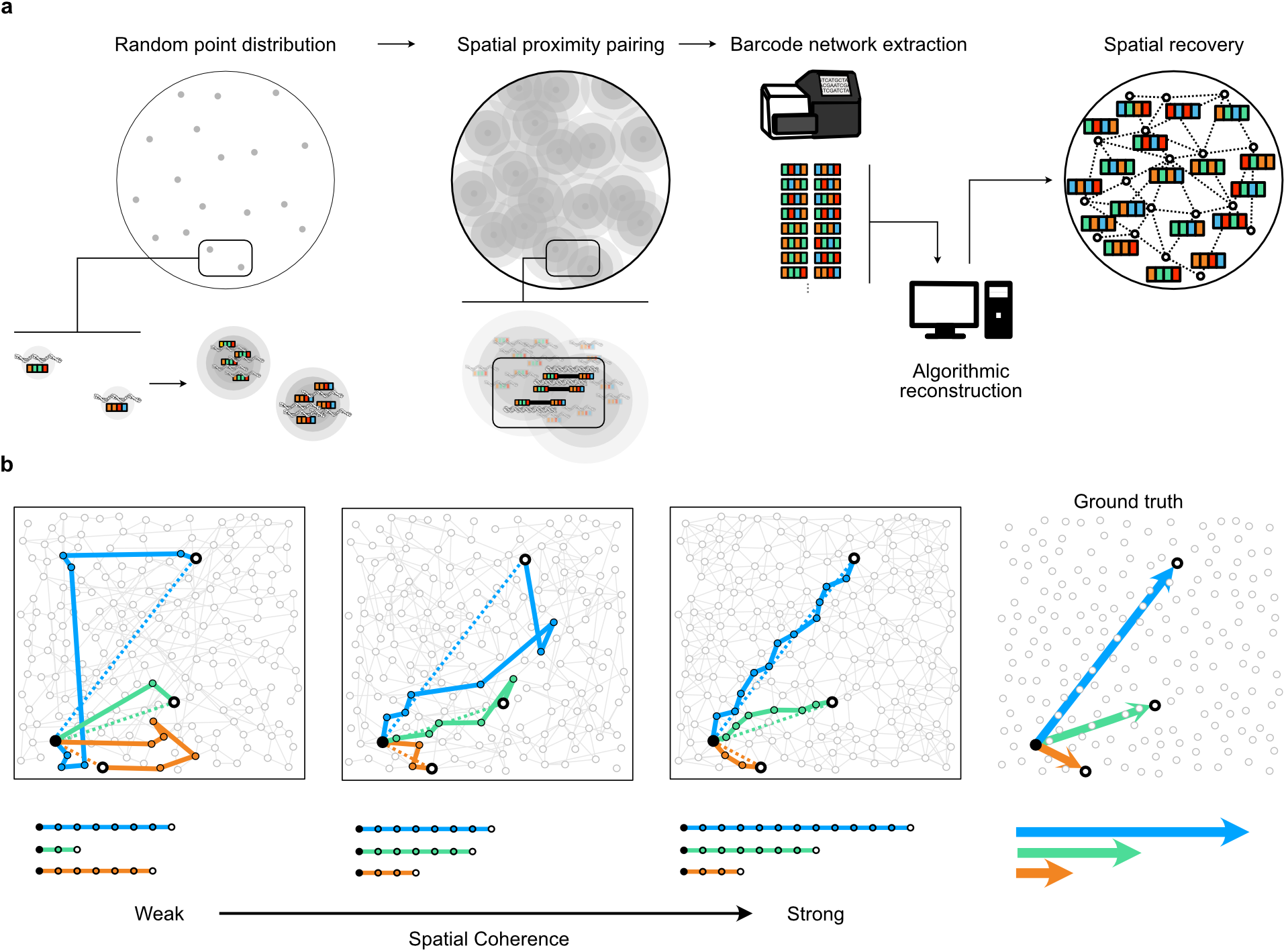
Spatial networks in sequencing-based microscopy. **a**. Sequencing-based microscopy pipeline schematic. DNA strands are distributed across space, each carrying a random DNA barcode. Seeded strands are clonally amplified forming polonies, each representing a region of space. Polonies link covalently, forming a proximity record. Strands are sequenced and converted to an edge list, and reconstructed into a spatial network whose structure preserves proximity relationships. **b** A progression of networks of increasing spatial coherence, showing the Euclidean distance ground truth (far right) and shortest path network hop distances for each topology. Networks with low spatial coherence have shortest path distances that correlate poorly to the ground truth Euclidean distances, versus strong correlation for spatially coherent networks.

Hence, a major challenge across this field is quality assessment without prior spatial position knowledge (ground truth). Quality metrics in simulation contexts can assess the quality of the reconstructions by direct comparison with ground truth (Bonet and Hoffecker, 2023), whereas laboratory experimentation demands secondary validation, e.g. observing known biological patterns of co-localization Karlsson et al. (2024), or comparing global morphology with expectations in model biological systems/organisms Qian and Weinstein (2023); Weinstein et al. (2019). These strategies depend on prior sample knowledge, are challenging to quantify, and may overlook distortions arising from false communication between uncorrelated network regions, sparse data, or sequencing/amplification errors.

A principles-based understanding of the structural network features that enable spatial information preservation would facilitate quality assessment without ground truth or prior knowledge, and aid progress in sequencing-based microscopy. We investigated the underlying mathematical relationship between network topology and spatial geometry, and identified a quantifiable property of spatially-informative networks. The fulfillment of this property, which we term “spatial coherence”, reflects the consistency with which a network’s relative topological distance relationships obey Euclidean geometric laws (Fig.1b). We show that spatial coherence is a feature of well-behaved spatial networks and conversely that the property is lacking in nonspatial and random networks and diminished when introducing random noise.

## Identifying geometric indicators of spatial networks

Whereas certain real-world networks exhibit a small-world property in which the shortest path between two nodes is short compared to network size (Watts and Strogatz, 1998; Barrat and Weigt, 2000), spatial networks, in contrast, exhibit a “large-world” property where traversing between distant points in physical space takes correspondingly many network hops in network space (Zemel and Carreira-Perpiñán, 2004). The network distance (known as shortest path distance) and the Euclidean distance are correlated (Hoffecker et al., 2019; Bonet and Hoffecker, 2023) (Fig.2a-c). For example, in the European high-speed rail network, it takes more connections to reach Rome from Stockholm than from Copenhagen. This correlation motivated us to interrogate classical geometric relationships while substituting Euclidean distance with network distance. Deviations from Euclidean behavior in these relationships offer a means to discriminate spatial and non-spatial networks without ground truth or prior sample knowledge.

**Figure 2.**
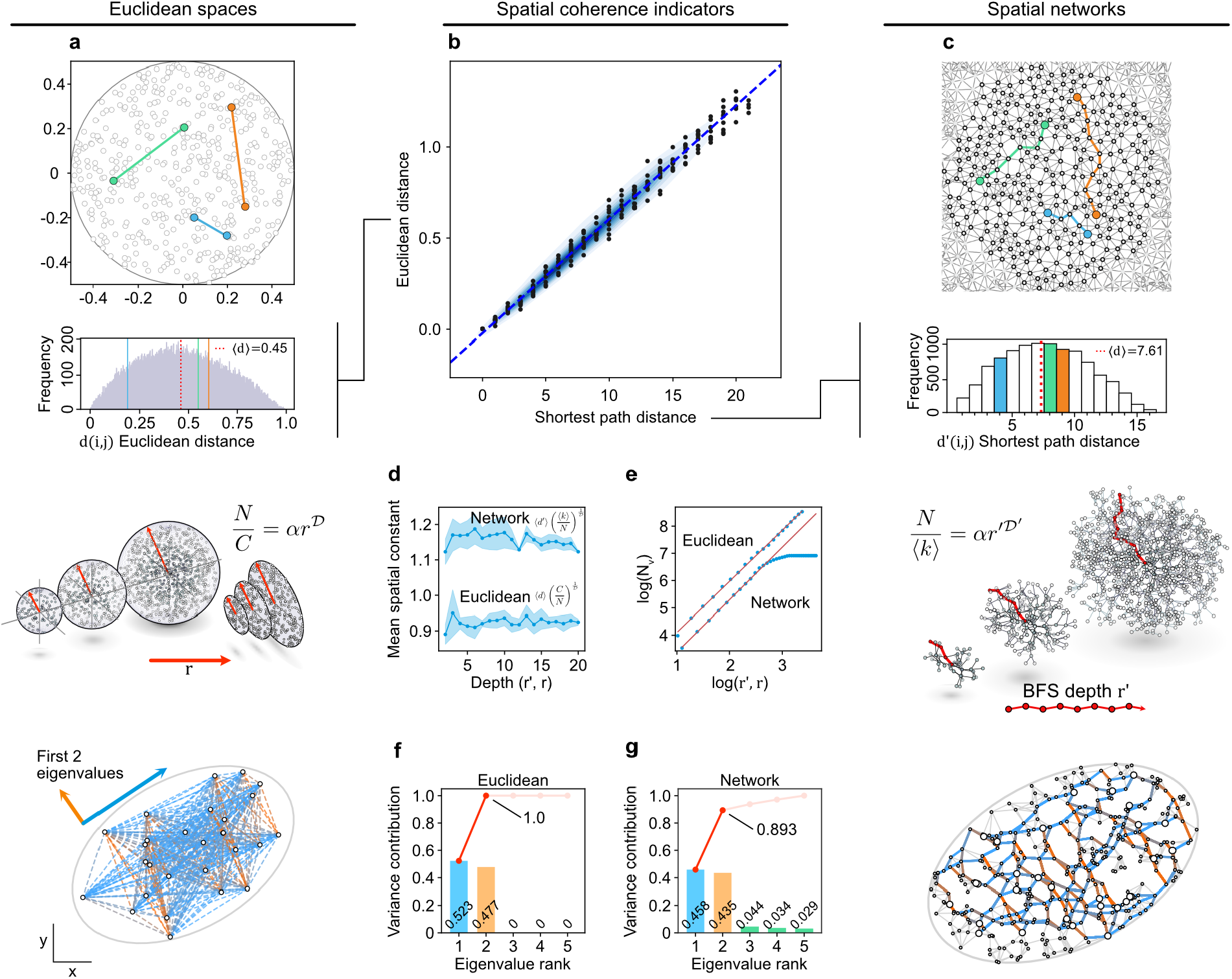
Geometric relationships for quantifying network spatial coherence. **a**. Selected pairwise Euclidean distances between points on a disc and their frequency. **b**. Network shortest path distances and Euclidean distances are highly correlated in spatial networks. **c**. A spatial network is composed of edges forming paths between nodes, and the shortest paths are analogous to Euclidean pairwise distances, with a corresponding frequency distribution. **d**. A plot of the dimensionless spatial constant, which relates pairwise Euclidean or shortest path distances to the volume of the space and should remain constant with space size. **e**. A log-log plot of the scaling law relating Euclidean or breadth-first-search radius to volume whose slope indicates intrinsic dimension of the space. **f**. The Gram matrix for a set of Euclidean points may be calculated from the pairwise distances of those points, the eigenvalues of which reveal the contributions to variation in the spread of points (left) from different dimensions (2 for spatially distributed points in 2D). **g**. The Gram matrix may also be computed from a set of shortest path distances substituting for Euclidean distances in the case of a spatial network. The first eigenvalues, representing the number of dimensions of the space, should account for the variation in distances (right) in a spatially coherent network. Contributions from higher dimensions indicate variation that cannot be explained by spatial dimensions.

We first sought to quantify the intrinsic dimension of a network via an analogy to Euclidean geometric scaling laws (Fig.2d,e), and to compare that dimension with the expected Euclidean dimension. The relationship between the number of points within a 𝒟-dimensional ball and its radius *r* follows a power law:

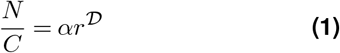

where *N* is the number of points, *C* is the concentration or point density, and *α* is a geometric factor (*α* = *π* for a circle, 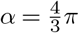 for a sphere). Similarly, the dimension of a network can be quantified using an analogous power law Shanker (2007), where *N* represents the number of moles, *C* the solution concentration, and the intrinsic network dimension 𝒟′ determined by fitting:

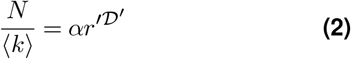

where the Euclidean radius *r* in Eq.1 has been substituted with the network distance *r*′ from a given origin node (i.e. the depth of a breadth-first-search, BFS), *N* is the number of nodes in the subgraph contained within *r*′ of a given origin node, ⟨*k*⟩ is the average node degree, *α* is a geometric factor. Here, spatial coherence would be indicated by a linear log *N* versus log *r*′ relationship and network dimension matching the dimensions of the physical space where the network formed.

Another quantity we examined is the consistency of dimensionless constants in geometric scaling laws, such as the relationship between the size of the space and the average distance between points. In Euclidean space, this expected distance ⟨*d*⟩ is related to the space size and the dimension 𝒟 by a constant:

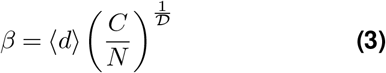

where 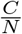 represents the size of the space, equivalent to volume in 3D or area in 2D. This constant remains consistent across scales in Euclidean space for a given dimensionality and boundary geometry (e.g., a cylinder vs. a sphere). The network-based analog of this dimensionless spatial constant is:

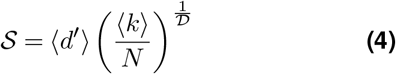

where *N* is the number of nodes, ⟨*d*⟩ has been substituted with the mean network distance ⟨*d*′⟩, and the point density (molar concentration) has been substituted with the average node degree ⟨*k*⟩. This constant may be computed either locally or as an ensemble property (Eq.4). A Euclidean spatial constant does not change across scales, and the extent to which this dimensionless quantity remains constant in a network would thus be an indicator of spatial coherence.

Finally, we explored a modified principal component analysis (PCA) approach for spatial network data. We analyzed the proportion of Gram matrix eigenvalues to determine the contribution of variation along each expected spatial dimension, substituting Euclidean with network distances Hastie et al. (2009) (Fig.2f,g). In Euclidean space, the first 𝒟 components account for all the variance, while the remaining dimensions do not contribute. Network distances whose first 𝒟 Gram matrix eigenvalues represent a proportion of the eigenvalues close to 1 would thus represent high spatial coherence, i.e. consistency with Euclidean space constraints.

### Proximity networks follow Euclidean geometry

We assessed the spatial coherence of multiple classes of well-behaved networks (Fig.3a). These networks, free of noise and artifacts, should all exhibit high spatial coherence. Except for a network of US county borders, a series of 2D, 3D, unipartite, and bipartite networks were simulated to capture a diverse range of architectures.

**Figure 3.**
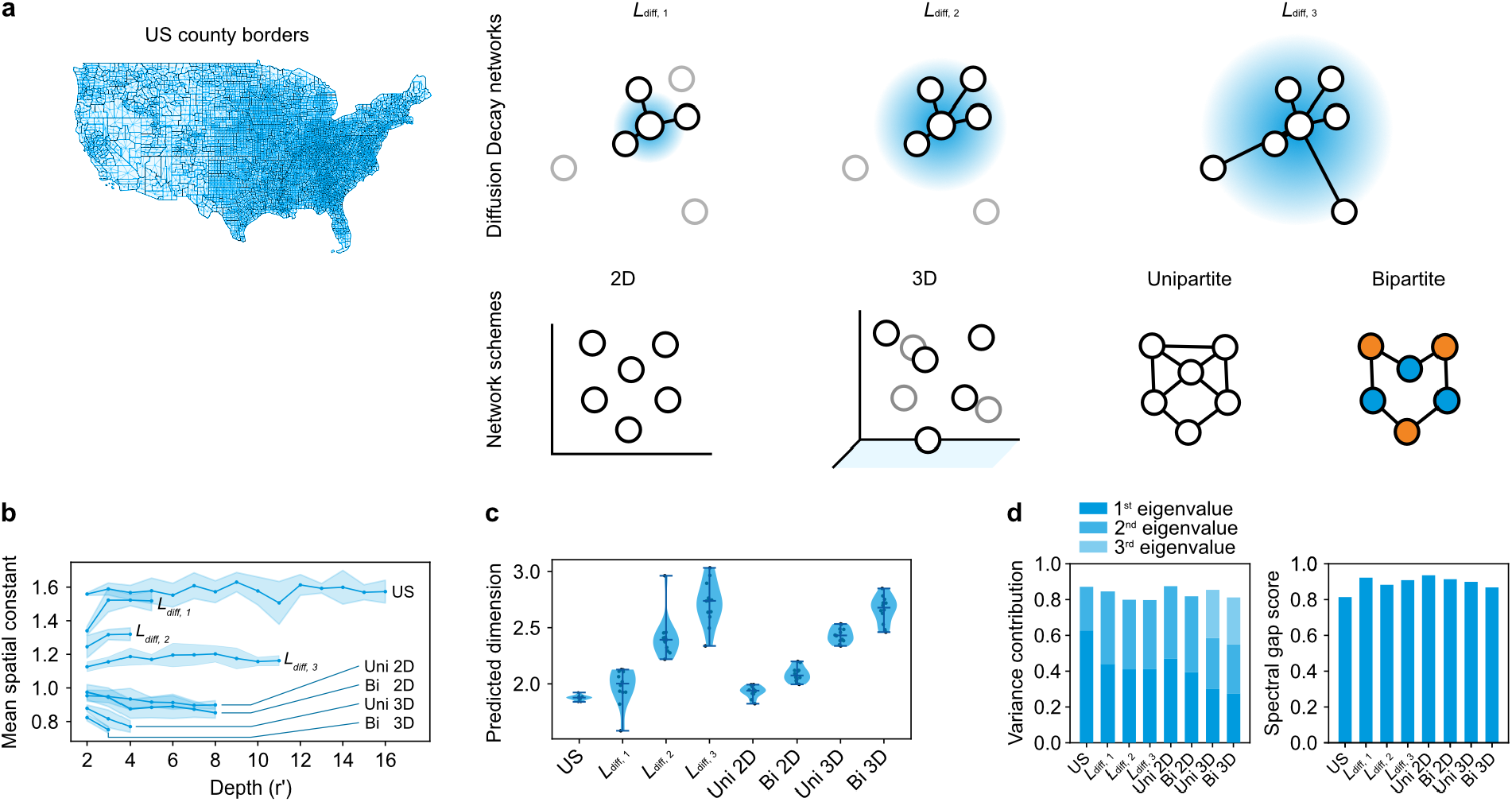
Spatial coherence for different network modalities. **a**. The different network modalities, including 2D/3D networks, unipartite/bipartite networks, and a distance decay based on diffusion to model interaction probabilities. Additionally, the US county borders network is included as a case study. **b**. Spatial constant against depth revealing stable profiles for the spatial networks examined. **c**. Network dimension values and **d**. spectral analysis of the Gram matrix: variance contribution and spectral gap score showing values for spatial networks approaching 1.

Using an exponential diffusion model where the probability of edge formation between two nodes is a function of their separation distance, we examined the impact of varied diffusion length *L*_diff_. The series of diffusion lengths that connect progressively distant parts of the network captures varying degrees of spatio-topological dependence, from strictly local to global network-spanning connectivity. The US county border network serves as a real network case study, exhibiting a non-uniform degree distribution, density variation, and non-isotropic boundary shape.

Spatial coherence measures applied to each network class (Fig.3b-d) exhibit a strong agreement with that which is expected of Euclidean data, despite fundamental architectural differences. Notably, the spatial constant exhibits a stable profile across the networks, despite variations in network depth. This is particularly clear in networks with high diffusion lengths and 3-dimensional networks, all nodes requiring fewer hops to reach one another.

The network dimensionality of US and unipartite 2D networks was near 2, while the bipartite 2D network and three-dimensional networks displayed a dimension between 2 and 3. Additionally, networks modeled with the decay function show increased variability in dimensionality, which could be attributed to the model’s stochasticity. In this progression, the network dimension increased from below 2 at the shortest diffusion length to approximately 3 at the maximum, suggesting that the perceived network dimensionality is sensitive even to non-random long-range connections.

Analysis of the Gram matrix eigenvalues shows variance contributions close to 1 by the primary dimensions for all networks. The US network stands out as its first eigenvalue dominates over the second. This may be attributed to its original rectangular shape, with one dimension contributing more than the other. On top of stable spatial constant profiles and intrinsic dimensionality values aligned with expectations, the high spectral gap scores across all networks establish these strictly generated distance-to-topology networks as spatially coherent.

### False edges result in spatially incoherent networks

While spatial networks with no artifacts have a high spatial coherence, approaching those of Euclidean spaces, real DNA barcode network data can contain false (nonspatially correlated) edges, shortcuts connecting distant spatial regions, e.g. via non-uniform polony growth or inconsistent diffusion, PCR errors introducing nonspatially correlated connections, or barcode collisions resulting in indistinguishable identities. Such deviations worsen image reconstruction. An open challenge in sequencing-based microscopy is to detect and quantify false edges, while the unavailability of ground truth prevents directly determining edges’ spatial correlation. We therefore sought to detect them with the spatial coherence framework relying purely on network data.

The increasing addition of randomly generated false edges to an otherwise spatially coherent network (Fig.4a) results in decreased correlation between Euclidean and network distances. This can be further visualized through the central node’s network distances which exhibit a radial progression in a clean spatial network that is replaced by a random pattern as false edges are added. These artifacts influence the quality of images(Fig.4b) reconstructed using the multiDimensional scaling algorithm. Here, false edges introduce conflicting distance constraints that require higher dimensions to resolve, analogous to the folding of a sheet in 3D.

**Figure 4.**
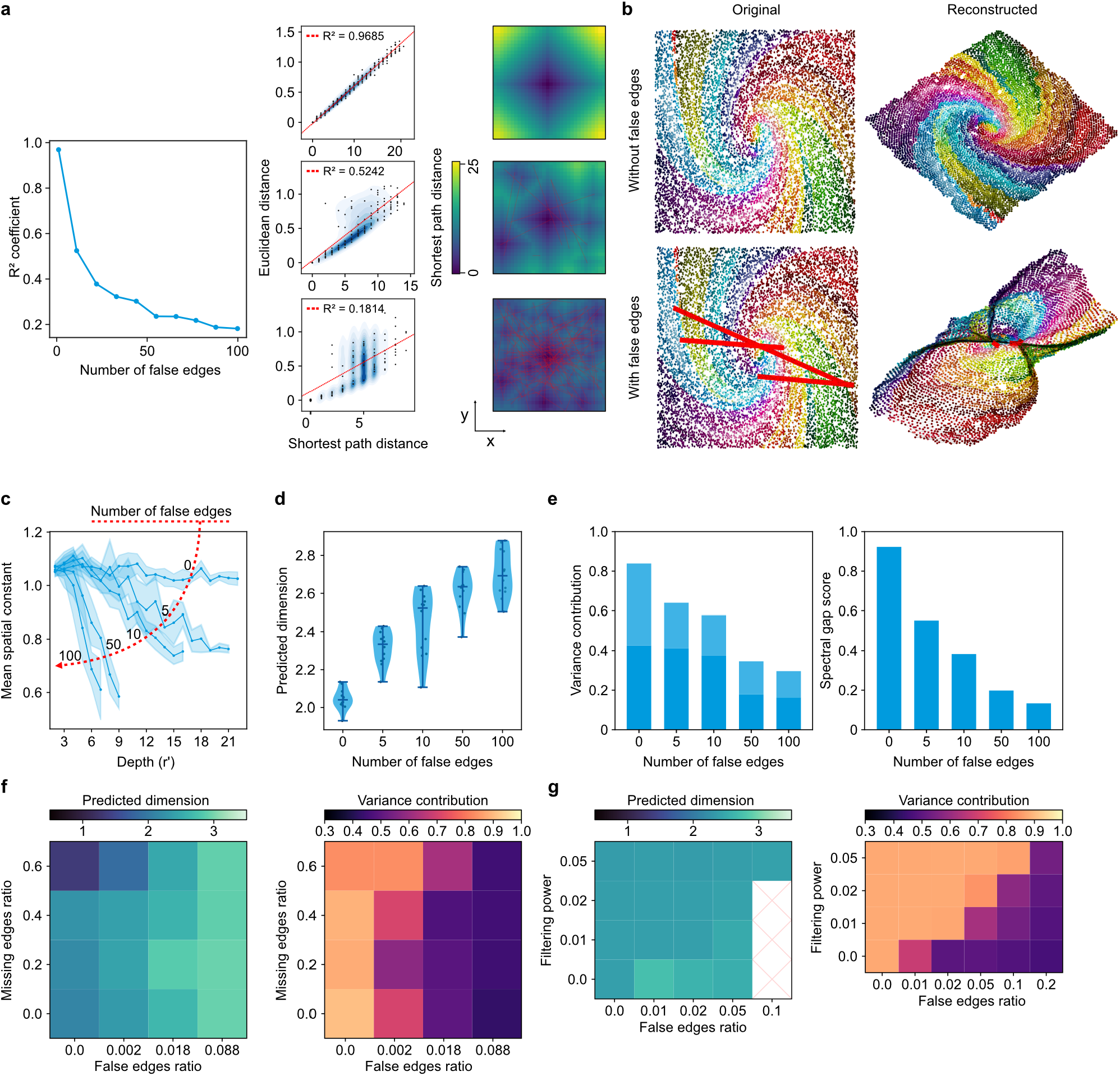
Impact of false edges in networks. **a**. Coefficient of determination *R*^2^ showing the correlation between Euclidean and shortest path distance for an initially spatially coherent network across different levels of false edges. Three samples selected to visualize such correlation, along with the shortest path distance from the central node. Behavior of the coefficient of determination *R*^2^ against the number of false edges introduced. **b**. Ground truth original positions colored according to a spiral pattern and the reconstructed positions using network information only, followed by the same data with added false edges (highlighted in red) connecting distant regions and its corresponding reconstruction. Spatial coherence measures in response to progressively more false edges: **c**. spatial constant values against BFS depth, **d**. network dimension estimates, and **e**. the variance contribution of the first 𝒟 eigenvalues along and spectral gap scores. **Effect of missing edges and error correction filtering on spatial coherence measures: f**. Dimension and Gram matrix variance contributions visualized across different false and missing edge frequencies. **g**. Dimension and Gram matrix variance contributions visualized for different false edge ratios and filtering powers.

We next sought to quantify the amount of false edges through the spatial coherence framework (Fig.4c-e). As false edges are progressively added to a well-behaved spatial network, the otherwise stable spatial constant profile drops, the network dimension increases from 2 to values near 3 -similar to a circular 1-dimensional ring with false edges being close to a 2-dimensional circle (Shanker, 2007)- and the spectral gap score decreases away from 1.

Spatial coherence is also impacted by missing edges, where points without an edge have a separation distance that would be compatible with an edge. Unlike false edges that connect distant regions, missing edges could emerge from insufficient interactions or shallow sequencing, with sparseness reducing the spatial constraints necessary for accurate network reconstruction. Because experimental data could potentially have both false and missing edges, we investigated their combined impact on spatial coherence (Fig.4f). False edges generally increase the network’s dimension and reduce the Gram matrix variance contribution, while missing edges lower the network dimension. In networks with 60% missing edges, the dimensionality approached 1. Notably, there is an interplay between false and missing edges that show a compensatory effect: the right proportion of false and missing edges makes the networks seem more spatially coherent, with a dimension closer to two and higher variance contributions. However, this effect was only seen when the false edges ratio is low. We then studied spatial coherence-based evaluation of filtering algorithms designed to remove false edges or fused nodes. Spatial coherence measures could help optimize filtering levels where ground truth is not known, thus avoiding the deletion of true edges while removing most false edges. We applied an error correction algorithm to simulated data Kloosterman et al. (2024), measuring the spatial coherence at different filtering powers (Fig.4g). Increasing filtering power enhanced spatial coherence, showing a network dimension closer to two and improving the Gram matrix variance contribution. A lower false edge ratio necessitated less filtering to maintain coherence, while higher ratios required more stringent filtering to achieve spatial coherence. This highlights the utility of spatial coherence as a criterion for selecting an appropriate filtering level for network data, reducing unnecessary pruning of useful data without the burden of extensive validation.

### Spatial coherence in sequencing-based microscopy datasets

We applied the spatial coherence framework to 2 published experimental datasets: Molecular Pixelation Karlsson et al. (2024) and DNA Microscopy Weinstein et al. (2019). Both datasets are 2D bipartite networks but differ in methodologies and target molecules. The Molecular Pixelation dataset focuses on the cell surface proteome, using rolling circle amplification to form DNA pixels capturing protein proximity. Nodes are referred to as pixels, and edges represent unweighted proximity interactions between protein antibodies and pixel products. DNA Microscopy targets the transcriptome with a diffusion-based method to generate DNA polonies (nodes), weighting edges by interaction frequency to form weighted networks. Weights were modeled using Fick’s laws to estimate physical distances, Methods C.4). Both are compatible with the spatial coherence framework, despite differences in data weighting and biological targets: shortest path distances for unweighted Molecular Pixelation and weight-to-distance transformation for the weighted DNA Microscopy.

We applied the framework to the Molecular Pixelation human PBMC dataset. We found that the spatial coherence framework was useful to efficiently benchmark and compare the hundreds of networks in the dataset. We used the Gram matrix eigenvalues (Fig.5a) to select three representative networks that we reconstructed with the STRND algorithm and for which we computed spatial coherence measures. Among these, PBMC 1 exhibited the best performance in metrics and could be reconstructed with short edges well-correlated to proximity. PBMC 2 showed some longer edges, while PBMC 3 had mostly distant connections. These observations, potentially attributable to the specifics of the reconstruction method, were corroborated with the spatial coherence measurements (Fig.5b-d). A simulated network provided a reference with a stable spatial constant, a network dimension close to 2, and high scores for the variance contribution and spectral gap for the Gram matrix. Similarly, PBMC 1 displayed high spatial coherence in all measures, whereas PBMC 2 and PBMC 3 exhibited declining spatial constant profiles, variable network dimensions close to 3 and lower scores for the Gram matrix spectral analysis.

**Figure 5.**
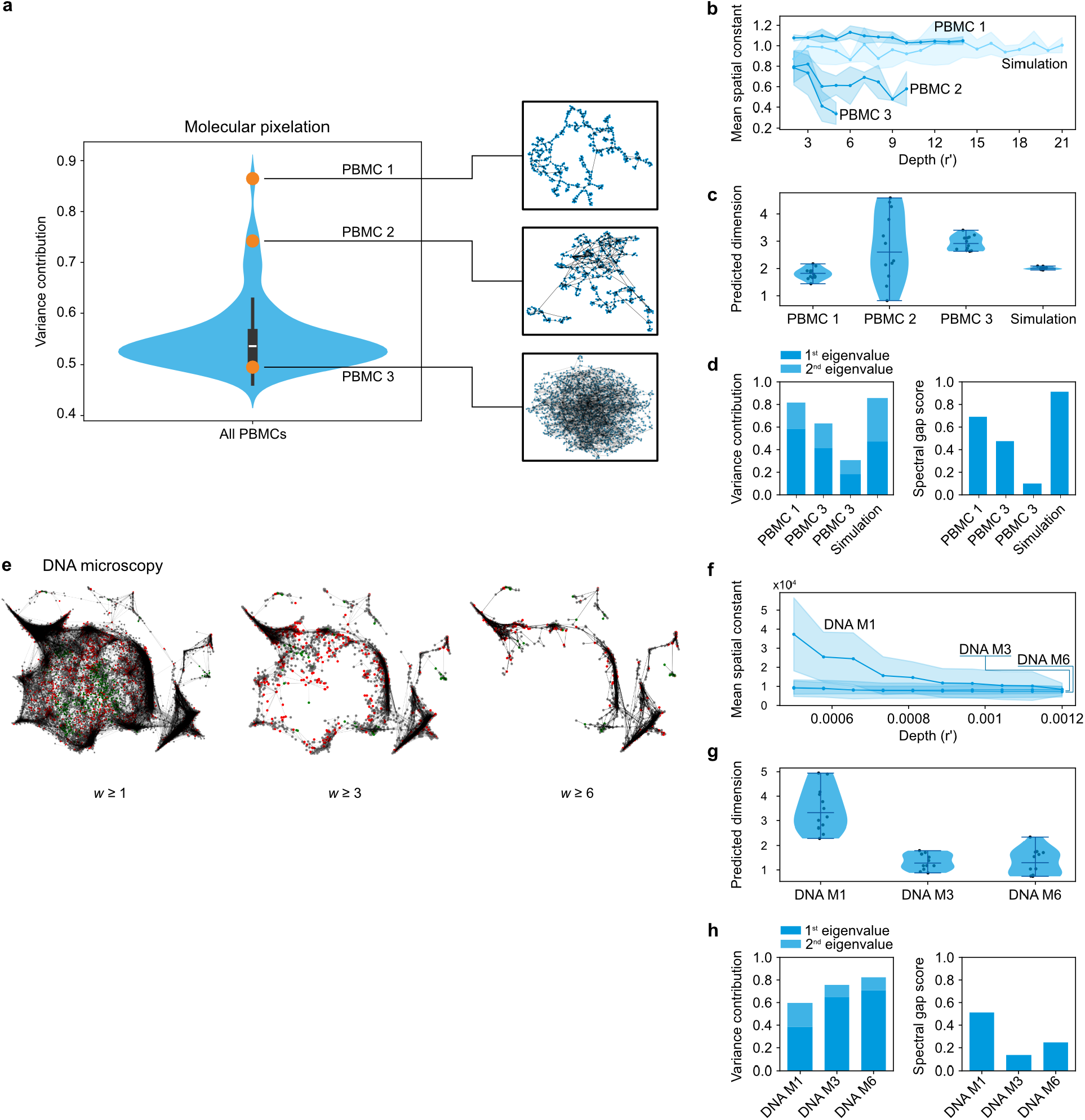
Spatial coherence measurement of published experimental data. **a**. Variance contribution of the Gram matrix eigenvalues for the PBMC dataset from Molecular Pixelation, where 3 representative networks are selected and reconstructed. Spatial coherence measures for these representative networks and comparison to an ideal simulated profile: **b**. Spatial constant profiles for the representative networks, **c**. predicted dimension measurements, and **d**. spectral analysis including variance contribution and spectral gap scores. **e**. Spatial coherence analysis of the DNA Microscopy dataset: Representative networks displayed using different weight filtering levels (with reconstructed locations taken from the sMLE result). Spatial coherence measures for different filtering levels of the DNA Microscopy dataset: **f**. spatial constant profiles, **g**. network dimension, and **h**. spectral analysis including variance contribution and spectral gap scores.

We analyzed the weighted DNA Microscopy network filtered by different thresholds of interaction weight (Fig.5e). We hypothesized that higher weights would be related to more reliable interactions, and found that it indeed improves spatial coherence (Fig.5f-h). The spatial constant stabilizes and the network dimension decreases to values below 2. The variance contribution of the Gram matrix eigenvalues also increases. However, higher filters lead to sparser networks, having a more 1-dimensional behavior. This is quantified both with the network dimension (values between 1 and 2) and the low contribution of the 2nd eigenvalue. The spectral gap score is also not optimal, with the 3rd largest eigenvalue of comparable magnitude to the 2nd.

## Conclusion

This study establishes the principle of spatial coherence as a fundamental property of spatial networks, demonstrating how it reflects the preservation of Euclidean geometric relationships when mapped onto network distances. By introducing this concept, we provide a way to analyze spatially informative networks without priors, particularly in the context of DNA barcode networks.

Our work identifies spatial coherence as a measurable property across both simulated and experimental networks and shows promise as a diagnostic tool for assessing data quality and detecting structural inconsistencies such as false edges. The three primary measures we developed — agreement between intrinsic network dimension and physical dimensions, stability of the spatial constant, and spectral analysis of the network distance Gram matrix — offer reliable metrics for evaluating spatial networks. Furthermore, these methods establish a common framework that can be applied across different technologies, improving comparability. Beyond immediate applications in validating the quality of networks, spatial coherence could serve as an optimization criterion for error correction. This correction can be improved by receiving feedback from spatial coherence measures, such as filtering power selection or custom loss functions. This approach could reduce the reliance on extensive prior knowledge or experimental validation, making high-confidence network analysis more accessible.

As spatial omics techniques evolve, the framework presented here can serve as a benchmark for evaluating and improving DNA barcode networks. By providing a common standard for assessing sequencing-based microscopy techniques, we hope to enhance the reproducibility and quality of spatial network data. This can represent a step forward to more accurate and reliable nanoscale, multiplexed imaging through DNA barcodes.

## Author Contributions

DFB implemented the algorithms, characterization, and computational exploration in the study. JIB and SL retrieved and analyzed data from the literature and run its algorithms. All authors contributed conceptual insights. DFB, ITH and SKD wrote the manuscript. DFB and ITH conceived the study.

## Competing Interests

All authors declare no competing interests

## Acknowledgements

We acknowledge support from the Swedish Research Council (no. 2020-05368 to ITH), and the European Research Council ERC (no. 949624 to ITH). The authors wish to thank Erik Benson (Karolinska Institutet), Antti Elonen and Pekka Orponen (Aalto University), and Ragnar Thobaben (KTH Royal Institute of Technology) for helpful discussions and insights.

## Methods

### A. Algorithm description

#### A.1. Spectral Analysis of the Gram Matrix

The rank of a Gram matrix derived from the pairwise distances of a set of points *X* is indicative of the dimension 𝒟 of *X*. Therefore, the first-𝒟 eigenvalues of the Gram matrix account for all the contribution while the rest of the spectra have null values. In particular, a Gram matrix is defined by the dot product of its set of points *X* as *G* = *XX*^*T*^. However, the point coordinates are not necessary to compute the Gram matrix as it is also related to the set’s pairwise distances *D* (Eq.5), known as the Euclidean Distance Matrix (EDM) Dokmanic et al. (2015):

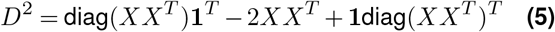

Here, **1** represents a column vector of all ones, and diag*X* denotes the column vector of the diagonal entries of the point coordinates matrix *X*.

This framework is generalized to networks, using the Shortest Path Distance Matrix *D*′ (SPDM) as opposed to the EDM *D*. To obtain the network’s Gram matrix *G* = *XX*^*T*^ Eq.5 is simplified by using the geometric centering matrix *J*

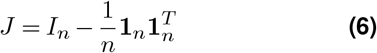

where *I*_*n*_ is the *n* × *n* identity matrix and **1**_*n*_ is an *n*-dimensional column vector of ones. Applying *J* results in positioning the centroid of *X* at the origin, effectively removing the terms diag(*XX*^*T*^)**1**_*T*_ and **1**diag(*XX*^*T*^)_*T*_ in Eq.5. This is known as double centering *D*^*′*2^. Therefore, the Gram Matrix can be obtained from *D*^*′*2^ by applying Eq.7:

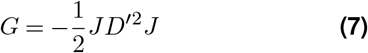

The eigenvalues of *G* coincide with the principal components obtained via Principal Component Analysis, up to a scaling factor Hastie et al. (2009). Therefore, the magnitude of the eigenvalues is linked with the variance explained by each eigenvector. The normalized contribution of the first 𝒟 eigenvalues (*λ*_1_, …, *λ*_*𝒟*_) is expressed as:

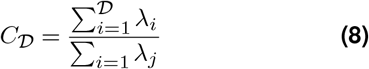

This measure is bounded between 0 and 1, and it reflects the extent to which the first 𝒟 dimensions capture the variance of the data.

An additional measure is that of the spectral gap, defined as the normalized difference between the last 𝒟 and the 𝒟 + 1 eigenvalue:

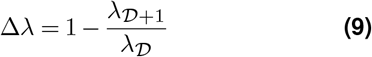

where Δ*λ* denotes the spectral gap. A larger gap indicates a greater distinction between the relevant dimensions and the residual dimensions, with the largest gap occurring in a case where the residual dimensions take null values.

#### A.2. Network Dimension

The network’s dimension is computed based on the Euclidean principle that the number of points (nodes) within a certain distance *r* increases according to a power-law, where the exponent corresponds to the dimension of the space. To estimate the network’s dimension, the shortest path distances *d*′ act as proxies for Euclidean distances *d*. In a large spatially coherent network, the network dimension should agree with the Euclidean dimension of the space. However, boundary effects influence the measurement, making it dependent on the origin point and the depth of the scaling relationship interrogated at that point. If the measurement begins near a boundary, the perceived dimensionality will be lower compared to a central start.

Therefore, several central network nodes are identified in order to minimize finite-size effects. Centrality in our network analysis is defined by the closeness centrality measure, which identifies nodes that are in proximity to all other nodes in the network, thereby minimizing finite-size effects. Closeness centrality for a given node is typically calculated as the reciprocal of the sum of the shortest path distances from that node to all other nodes in the network. Mathematically, for a node *i*, the closeness centrality *CC*_*i*_ is defined as:

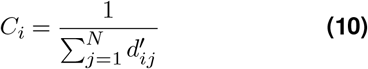

where 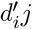 is the shortest path distance between node *i* and node *j*, and *N* is the total number of nodes in the network. Nodes with higher closeness centrality are used as starting points for a BFS of depth *r*′, as they are statistically more likely to represent the central structure of the network and reduce boundary biases in network traversal.

After central nodes are identified, the number of nodes *N* within a depth *r*′ is computed for all possible values of *r*′, starting at *r*′ = 1, where *N* represents the number of immediate neighbors one hop away from the origin. The gathered data is used to compute the fit log(*N*) against log(*r*′) that enables finding the power defining the power-law behavior, or the network’s dimension. To ensure linearity, a sliding window of points is iteratively selected, and the segment providing the highest coefficient of determination is chosen. The slope of this linear fit is used to estimate the network’s dimension, which in the case of a proximity network closely aligns with the dimensionality of the underlying Euclidean space.

#### A.3. Spatial Constant

Analogous to the mean line segment in Euclidean space, the spatial constant 𝒮 measurement is concerned with the mean distance between all pairs of nodes.

In Euclidean space, the mean line segment of a circle is Solomon (1978):

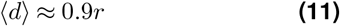

where *r* is the radius of the circle. But *N* = *ρV* = *ρπr*^2^, and Eq.11 becomes

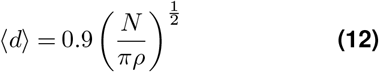

The spatial constant is chosen by definition as the dimensionless parameter 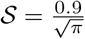 that does not depend on density nor the number of points. By rearranging the terms, it can be generalized to the network’s case:

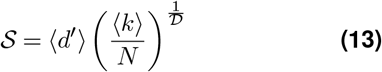

where ⟨*d*⟩ is the mean shortest path distance between nodes, *N* is the number of nodes, 𝒟 is the network’s original dimension, and ⟨*k*⟩ is the average degree of the network. The reason for using ⟨*k*⟩ as an analog of the Euclidean density *ρ* is because the average degree accounts for how many nodes there are on average at one space unit.

The assessment of the spatial constant’s behavior is done by sampling the network at different size scales. First, origin nodes are selected at random and a BFS traversal of fixed depth is conducted. This depth is increased iteratively, allowing for the discovery of more nodes and effectively increasing the size *N*. Different BFS depths represent different network sizes. Therefore, this approach measures how the spatial constant changes with network growth. A consistent spatial constant that mirrors the behavior in Euclidean spaces would suggest the absence of false edges in the network. Conversely, a varying spatial constant indicates that the network’s average shortest path length varies across different scales, pointing out that the behavior is not consistent and could potentially have false edges.

### B. Algorithm implementation

All algorithms are implemented in Python 3.10 and the code will be publically available both as a GitHub repository and as a Python package. The implementation relies at its core on efficient network representations, shortest path distance computations, and BFS traversals.

#### B.1. Network representation

This study takes advantage of sparse matrix representations for network data, specifically using the Compressed Sparse Row (CSR) format. The choice is motivated by the efficiency in storing and manipulating sparse graphs, which have only a fraction of the potential *N* (*N* −1)*/*2 edges. Spatial networks are sparse in that sense because the number of edges is usually bounded by spatial constraints and scales proportionally to the number of nodes *N* ∝ *E*. Using a CSR format can accelerate procedures to compute network properties, such as degree distribution and shortest path distance matrices. However, for the latter, they are memory intensive to store as the space requirements scale quadratically with the number of nodes.

#### B.2. Shortest Path Distance Matrix

The SPDM *D*′ denotes the shortest path distance *D*′_*ij*_ between nodes *i* and *j*. To compute the SPDM from the CSR representation of a network, we employ Dijkstra’s algorithm, optimized for sparse matrices. In particular for this work, the shortest path distance matrix of a network is used to obtain the eigenvalues of its Gram matrix and to compute the network’s dimension.

#### B.3. Breadth-First Search

Breadth-First Search (BFS) is implemented by selecting an origin node and using a queue to keep track of the next nodes to visit. The depth of the traversal can be chosen to study the network at different scales, where higher depths result in bigger scales. Exploring different scales is used for the spatial constant computation, where several origin nodes are selected and BFS traverals with varying depths are run from them. This results in multiple subgraphs representing different network regions, each with its corresponding spatial constant.

#### B.4. Large networks

In scenarios where the network’s scale makes computations such as the shortest path matrix or eigenvalue decompositions prohibitively expensive, especially in terms of space complexity, the network can be sampled. This is done by selecting a starting node and running a BFS traversal until a certain number of nodes is reached. This threshold is user-customizable and can be adapted depending on hardware.

### C. Simulated data processing

#### C.1 Synthetic data set

Networks are simulated by generating uniformly distributed points within a confined region, by default a circle in 2D and a sphere in 3D. The geometric shape of these distributions primarily affects the boundaries and has minimal impact on the bulk properties of the network. After generating points, networks are created based on the criterion that spatially close points form edges. Specifically, Fig.3 includes unipartite, bipartite, 2D, and 3D networks to model various experimental setups. Unipartite networks can form edges with all pairs of nodes, while bipartite networks consist of two distinct node sets where connections only occur between nodes from different sets. While existing research in sequencing-based microscopy predominantly uses 2D bipartite schemes Weinstein et al. (2019), Karlsson et al. (2023), our dataset also explores 3D networks due to growing interest in this modality Qian and Weinstein (2023). Unless otherwise stated, the K-Nearest Neighbor (KNN) proximity is used with *K* = 6 in 2D and *K* = 15 in 3D, based on the average degree in Voronoi tessellations. Bipartite networks establish connections using the K-nearest neighbors within their allowed node sets, ensuring no two nodes within the same set share an edge.

Furthermore, we also model connectivity via diffusion with an exponential distance decay in the interaction probability:

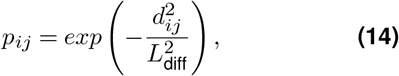

where *p*_*ij*_ is the probability of interaction, *d*_*ij*_ is the pairwise distance and *L*_diff_ is the diffusion length, or the characteristic length scale of the exponential decay. The larger the diffusion length, the more likely are interactions between far apart molecules.

For the specific experimental setting of DNA microscopy on diffusing amplicons Weinstein et al. (2019), the simulated data set is obtained by running the pipeline at https://doi.org/10.5281/zenodo.10256692 Kloosterman et al. (2024) with a default amplitude and spread of 10. The filtering algorithm from the same pipeline is used to study the spatial coherence before and after filtering.

#### C.2. False edges and missing edges

Modifications in the original network’s topology are introduced in the form of false edges and missing edges. False edges are created by selecting pairs of nodes that are not originally connected and linking them. This is done until the desired false edges number or ratio is satisfied. Conversely, missing edges are created by removing existing edges. If the deletion disconnects the network, another pair of nodes is chosen. As with false edges, the process is iterated until the desired sparsity of ratio is satisfied.

#### C.3. US County dataset

Data was acquired from https://census.gov/cgi-bin/geo/shapefiles/index.php?year#2023&layergroup=Counties+%28and+equivalent%29. The resulting shapefile was processed using Python’s package geopandas. The inferred network was created by connecting mainland counties based on adjacency.

#### C.4. Retrieval of published experimental data

The Molecular Pixelation PBMC dataset was downloaded from https://software.pixelgen.com/datasets/1k-human-pbmcs-v1.0-immunology-I and was processed according to the default guidelines https://software.pixelgen.com/mpx-analysis/tutorials/introduction. In particular, network components (putative cells) with 2000 or less edges were not accounted for and the metric *τ*, which measures the skewness of antibody counts, was used to identify outliers Yanai et al. (2005). The representative networks PBMC 1, PBMC 2, and PBMC 3 correspond to, respectively, the networks originally named RCVCMP0001392, RCVCMP0002024, and RCVCMP0000120. The DNA Microscopy dataset was retrieved from https://www.ncbi.nlm.nih.gov/sra via accession number PRJNA487001 and processed according to code from https://github.com/jaweinst/dnamic to perform read count filtration of 2, deduplication based on edit distance, and reconstructed into an image to reproduce the result of Weinstein et al.

##### Weighted Networks

To simulate a weighted experimental setup where weights correspond to polony interactions we follow the model proposed in previous studies Weinstein et al. (2019); Kloosterman et al. (2024) which is based on Fick’s law of diffusion. The model relates the reaction rate *L*_diff_ and the distance between two polonies, *i* and *j*, each of different types. This relationship is expressed by the equation:

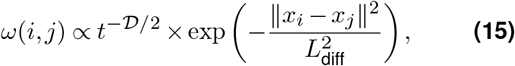

where *t* represents the time since the start of the information transference step, 𝒟 is the space dimensionality, and *x*_*i*_, *x*_*j*_ are the coordinates of polonies *i* and *j*, respectively. The characteristic diffusion length *L*_diff_ is defined as:

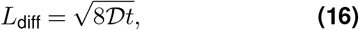

with *D* being the diffusion constant. The diffusion constant value strongly depends both on the medium where DNA is diffusing and the length of the DNA strands (citation). The variation can be several orders of magnitude. However, considering the known results for agarose gels (electrophoresis) with 2% agarose concentration and a DNA strand length of about 10_2_, *D* is roughly of the order 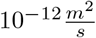. Therefore, if the DNA strands are diffusing for some minutes the order of magnitude of the characteristic length *L* will be about 10 micrometers.

Based on the diffusion model, the number of reactions between each pair of nodes can be estimated to determine the edge weights in the network

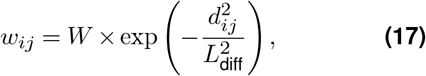

where *w*_*ij*_ denotes the edge weight between nodes *i* and *j, d*_*ij*_ its corresponding distance, *W* is the amplitude reflecting the inherent reactivity of the polonies and sequencing depth, and *L*_diff_ indicates the effective interaction range via diffusion. The amplitude affects the likelihood and intensity of reactions, thereby influencing the weight of the connections in the resulting network graph.

#### C.5. Impact of Diffusion Constant on Inferred Distances

Analyzing the relationship between the interaction range *L*_diff_ and the inferred distances, we derive the following from Eq. 17:

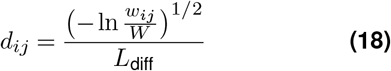

The equation indicates that changes in *L*_diff_ scale the computed distance *d*_*ij*_ without altering the relative distribution among different pair distances. Consequently, variations in *L*_diff_ act as scale modifiers and do not impact the normalized or relative spatial distribution of node distances. Therefore, the interaction range is not a critical parameter for inferring the topology of the network in this model. Its primary role is establishing the scale of measurement rather than influencing the actual spatial configuration. As such, the model primarily relies on the amplitude *W*, which determines the reaction rate and can be found experimentally from peak weight observations.

## Code Availability

The code containing the spatial coherence framework and additional tools to process data can be found at https://github.com/DavidFernandezBonet/Network_Spatial_Coherence.

## Extended Data

**Figure S1.**
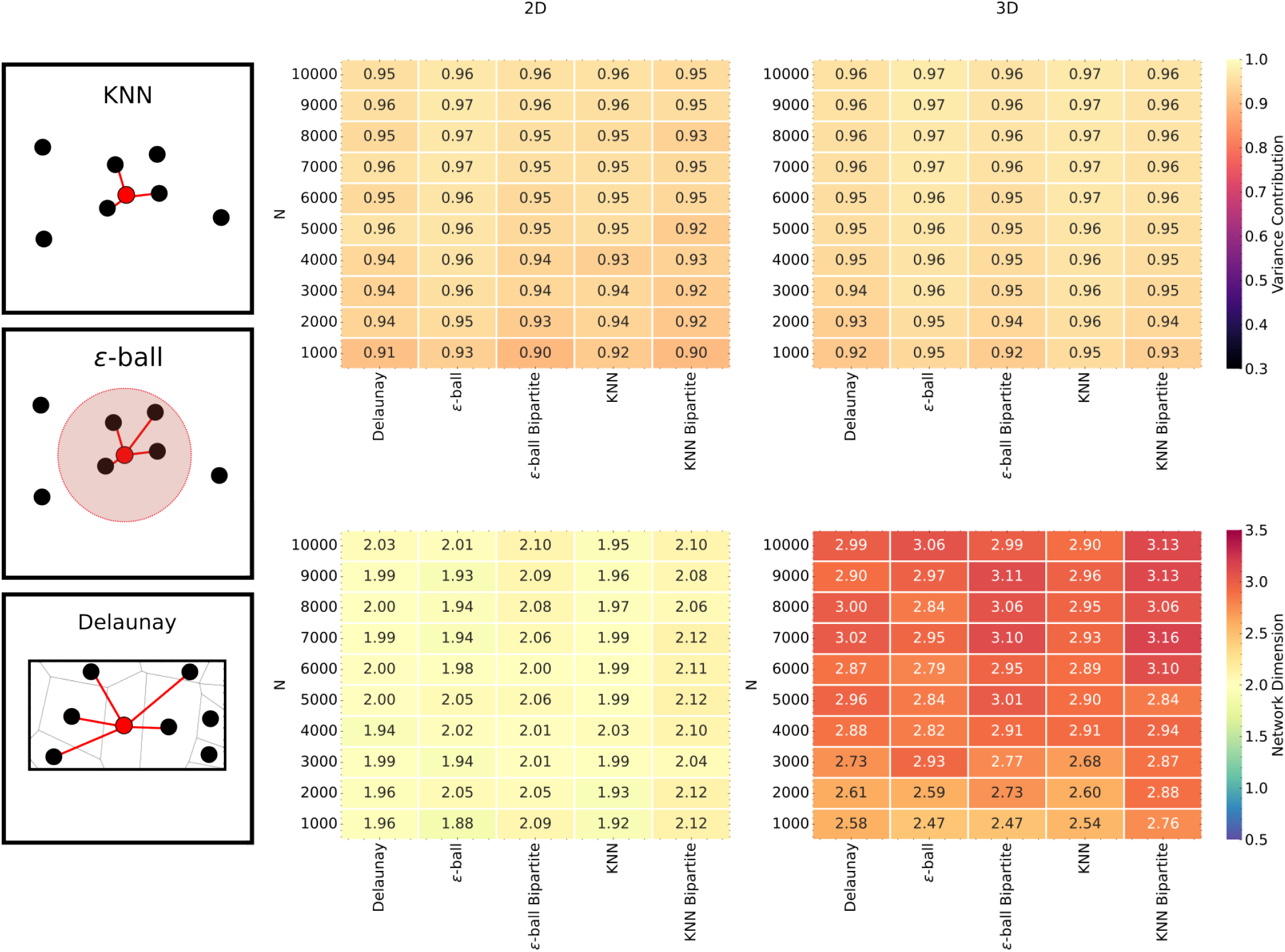
**Spatial coherence measures are analyzed across different numbers of nodes and proximity graph modalities in both 2D and 3D configurations. The proximity graph modalities examined include K-Nearest Neighbors (KNN)**, *ϵ***-ball, and Delaunay graphs. In the KNN modality, connections are formed between each node and its nearest K neighbors. In the** *ϵ***-ball modality, nodes are connected if they are within a radius** *ϵ* **of each other. In the Delaunay modality, connections are established between nodes with adjacent Delaunay cells. The bipartite case is also presented where applicable**.

**Figure S2.**
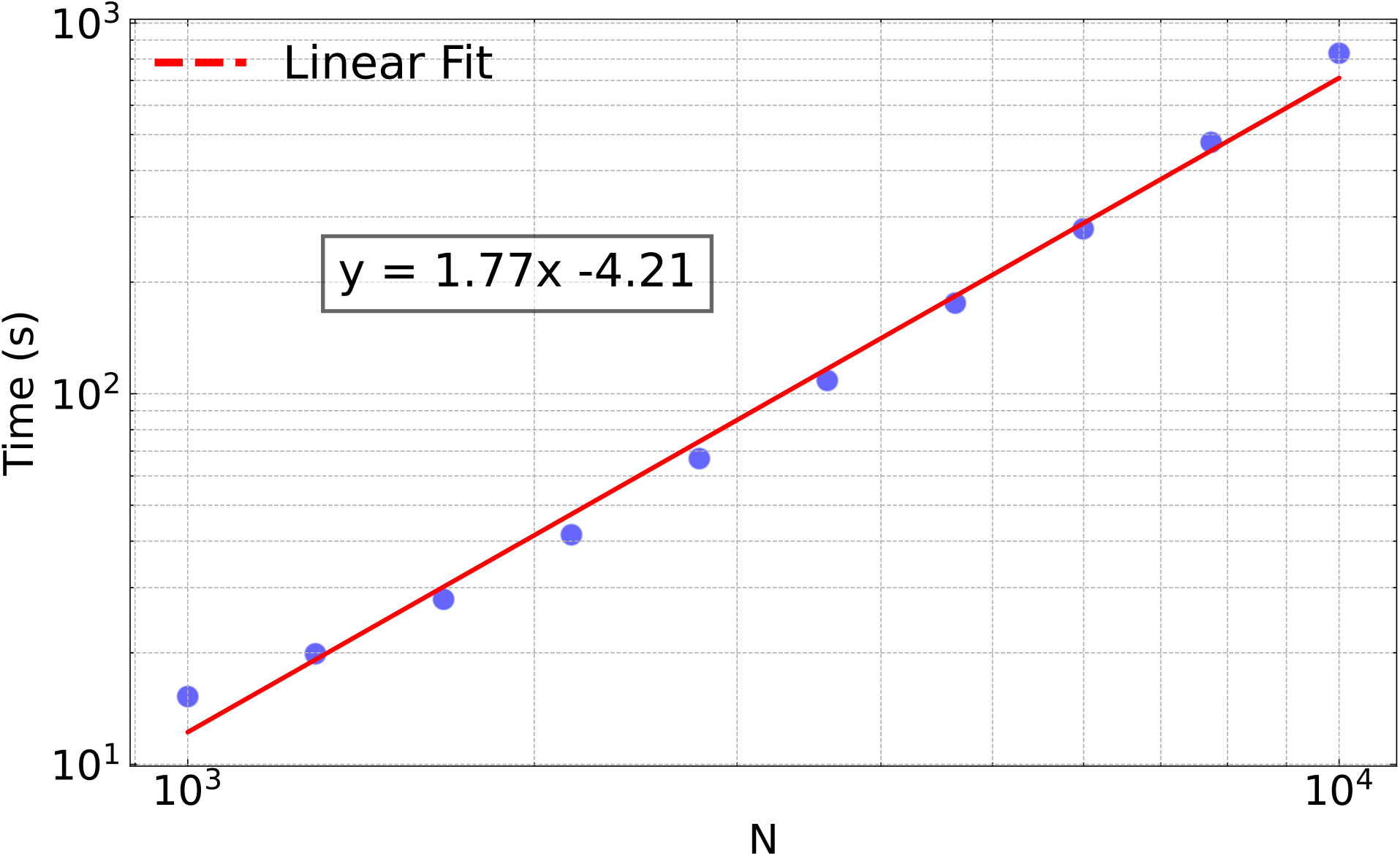
Empirical analysis of the time complexity associated with the spatial coherence pipeline. Computational time scales with varying input graph sizes, with a sub-quadratic scaling of 1.77.

**Figure S3.**
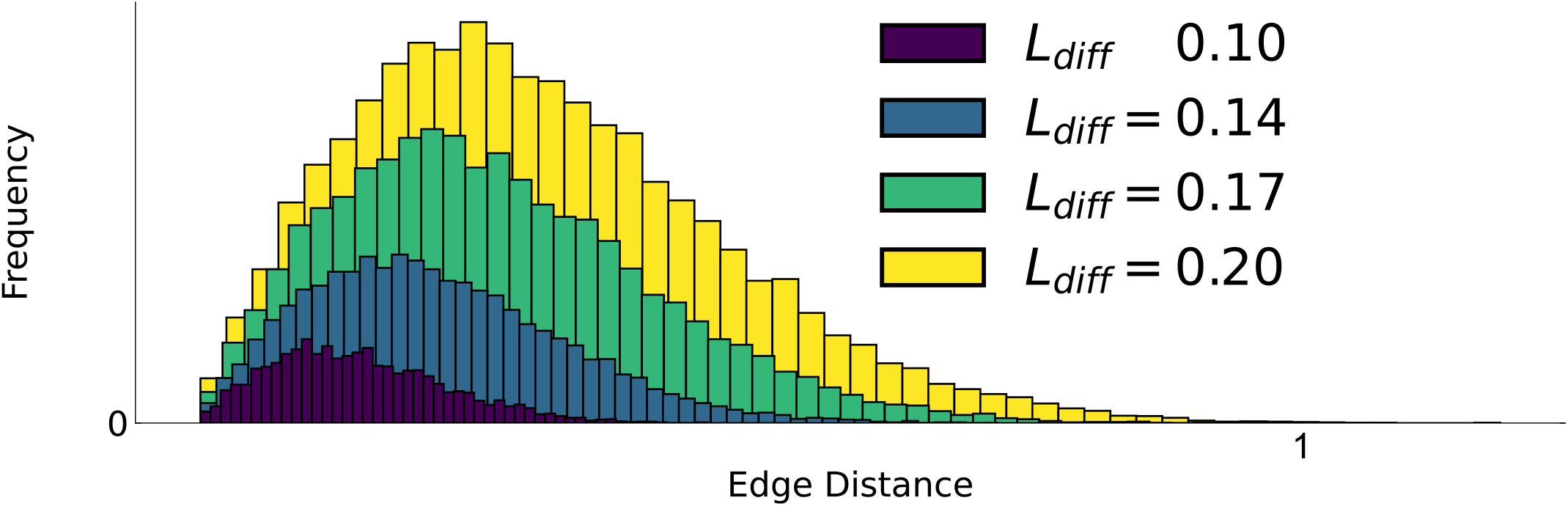
Edge length distribution for different characteristic diffusion lengths *L*_*diff*_ .

**Figure S4.**
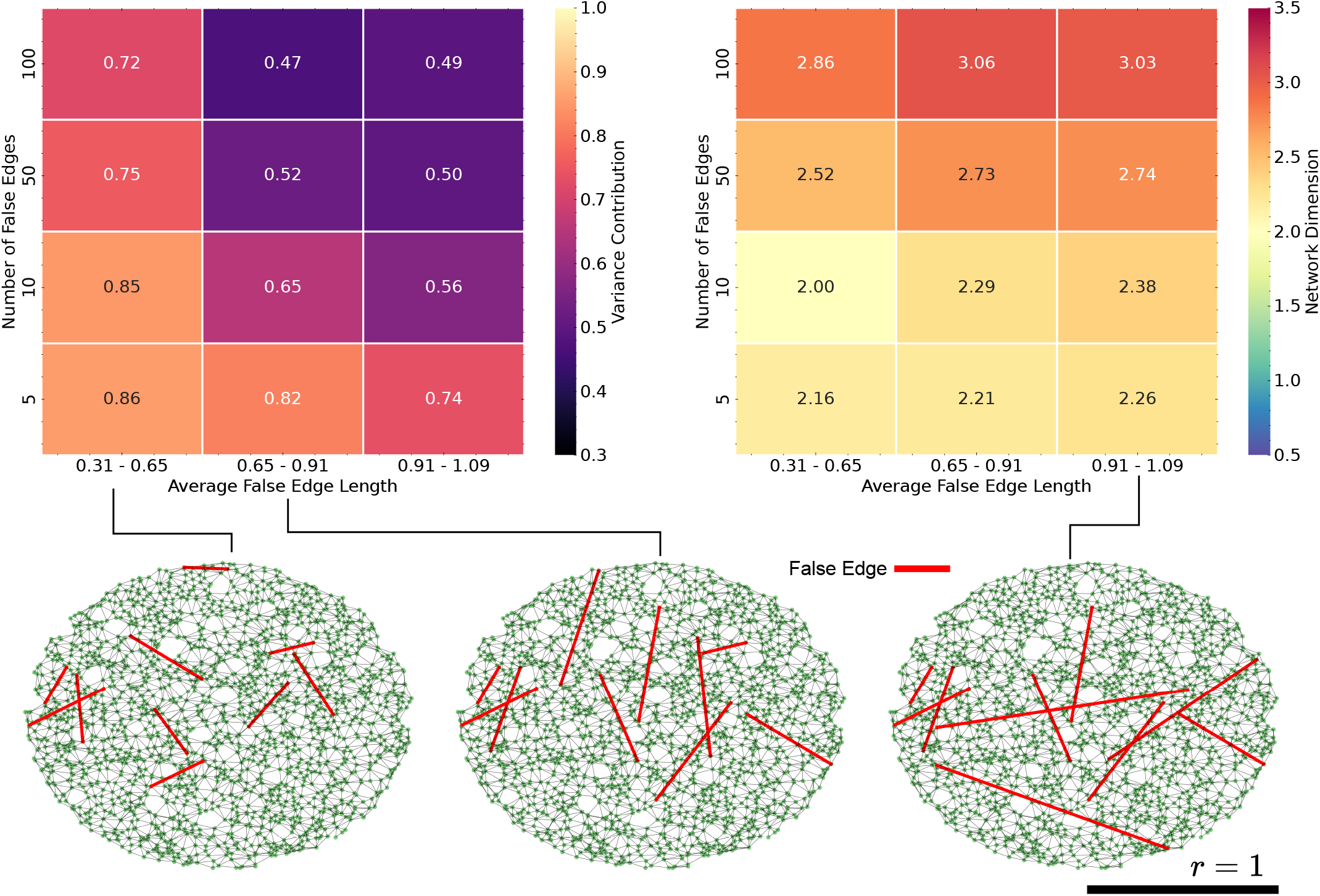
Impact of both the number and length of false edges on spatial coherence measures. An increase in length of false edges leads to a decrease in spatial coherence.

**Figure S5.**
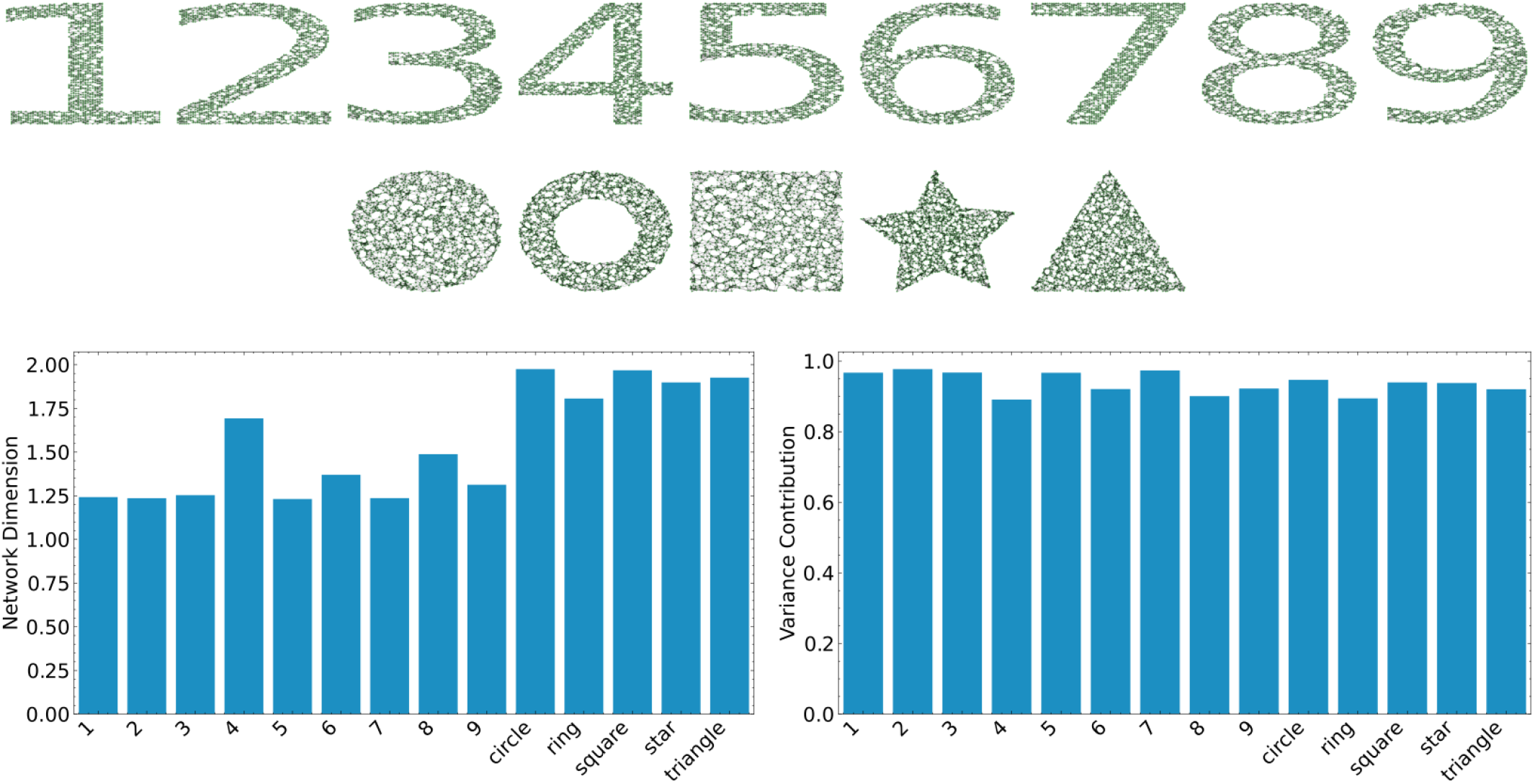
Exploration of different point cloud geometric configurations with different topologies and its effect on spatial coherence.

**Figure S6.**
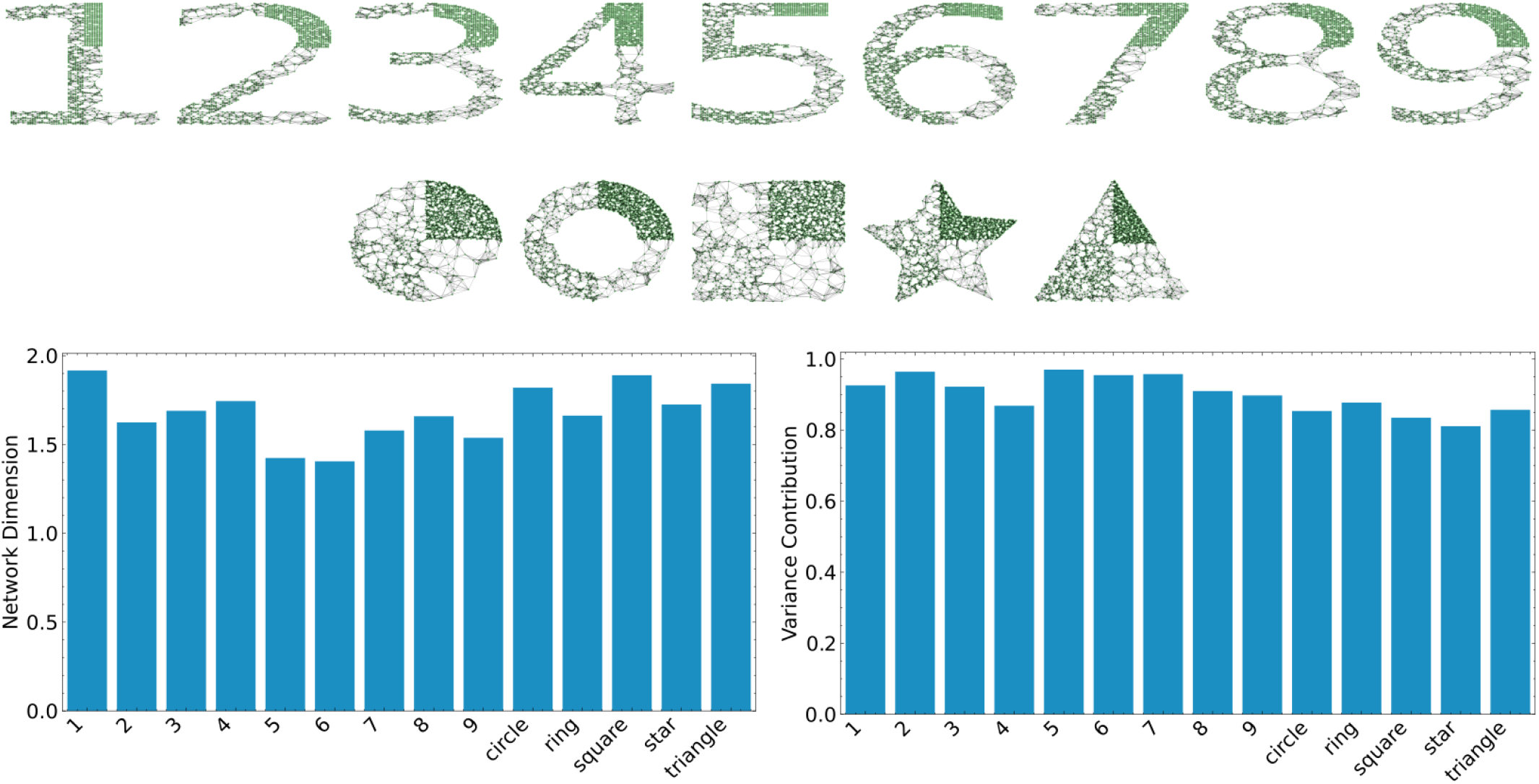
Effect of density anomalies in spatial coherence measures, featuring regions of varying density and shapes.

## Notes

### Competing Interest Statement

The authors have declared no competing interest.

### Summary of Updates

Figures have been updated and new supplementary figures added.

## Bibliography

Achim, K., Pettit, J.-B., Saraiva, L. R., Gavriouchkina, D., Larsson, T., Arendt, D., and Marioni, J. C. High-throughput spatial mapping of single-cell rna-seq data to tissue of origin. Nature biotechnology, 33(5):503–509, 2015.

Barrat, A. and Weigt, M. On the properties of small-world network models. The European Physical Journal B-Condensed Matter and Complex Systems, 13:547–560, 2000.

Bonet, D. F. and Hoffecker, I. T. Image recovery from unknown network mechanisms for dna sequencing-based microscopy. Nanoscale, 15(18):8153–8157, 2023.

Boulgakov, A. A., Xiong, E., Bhadra, S., Ellington, A. D., and Marcotte, E. M. From space to sequence and back again: Iterative dna proximity ligation and its applications to dna-based imaging. bioRxiv, page 470211, 2018.

Boulgakov, A. A., Ellington, A. D., and Marcotte, E. M. Bringing microscopy-by-sequencing into view. Trends in biotechnology, 38(2):154–162, 2020.

Dokmanic, I., Parhizkar, R., Ranieri, J., and Vetterli, M. Euclidean distance matrices: essen-tial theory, algorithms, and applications. IEEE Signal Processing Magazine, 32(6):12–30, 2015.

Gawad, C., Koh, W., and Quake, S. R. Single-cell genome sequencing: current state of the science. Nature Reviews Genetics, 17(3):175–188, 2016.

Glaser, J. I., Zamft, B. M., Church, G. M., and Kording, K. P. Puzzle imaging: Using large-scale dimensionality reduction algorithms for localization. PloS one, 10(7):e0131593, 2015.

Gopalkrishnan, N., Punthambaker, S., Schaus, T. E., Church, G. M., and Yin, P. A dna nanoscope that identifies and precisely localizes over a hundred unique molecular fea-tures with nanometer accuracy. bioRxiv, 2020.

Greenstreet, L., Afanassiev, A., Kijima, Y., Heitz, M., Ishiguro, S., King, S., Yachie, N., and Schiebinger, G. The dna-based global positioning system—a theoretical framework for large-scale spatial genomics. bioRxiv, 2022.

Halpern, K. B., Shenhav, R., Matcovitch-Natan, O., Toth, B., Lemze, D., Golan, M., Massasa, E. E., Baydatch, S., Landen, S., Moor, A. E., et al. Single-cell spatial reconstruction reveals global division of labour in the mammalian liver. Nature, 542(7641):352–356, 2017.

Hastie, T., Tibshirani, R., and Friedman, J. H. The Elements of Statistical Learning: Data Mining, Inference, and Prediction, volume 2. Springer, 2009.

Hoffecker, I. T., Yang, Y., Bernardinelli, G., Orponen, P., and Högberg, B. A computational framework for dna sequencing microscopy. Proceedings of the National Academy of Sciences, 116(39):19282–19287, 2019.

Karaiskos, N., Wahle, P., Alles, J., Boltengagen, A., Ayoub, S., Kipar, C., Kocks, C., Rajewsky, N., and Zinzen, R. P. The drosophila embryo at single-cell transcriptome resolution. Science, 358(6360):194–199, 2017.

Karlsson, F., Kallas, T., Thiagarajan, D., Karlsson, M., Schweitzer, M., Navarro, J. F., Leijonancker, L., Geny, S., Pettersson, E., Rhomberg-Kauert, J., et al. Molecular pixelation: Single cell spatial proteomics by sequencing. bioRxiv, pages 2023–06, 2023.

Karlsson, F., Kallas, T., Thiagarajan, D., Karlsson, M., Schweitzer, M., Navarro, J. F., Leijonancker, L., Geny, S., Pettersson, E., Rhomberg-Kauert, J., et al. Molecular pixelation: spatial proteomics of single cells by sequencing. Nature Methods, pages 1–9, 2024.

Ke, R., Mignardi, M., Pacureanu, A., Svedlund, J., Botling, J., Wählby, C., and Nilsson, M. In situ sequencing for rna analysis in preserved tissue and cells. Nature methods, 10(9): 857–860, 2013.

Kloosterman, A., Baars, I., and Högberg, B. An error correction strategy for image reconstruction by dna sequencing microscopy. Nature Computational Science, pages 1–9, 2024.

Lee, J. H., Daugharthy, E. R., Scheiman, J., Kalhor, R., Ferrante, T. C., Terry, R., Turczyk, B. M., Yang, J. L., Lee, H. S., Aach, J., et al. Fluorescent in situ sequencing (fisseq) of rna for gene expression profiling in intact cells and tissues. Nature protocols, 10(3):442–458, 2015.

Lewis, S. M., Asselin-Labat, M.-L., Nguyen, Q., Berthelet, J., Tan, X., Wimmer, V. C., Merino, D., Rogers, K. L., and Naik, S. H. Spatial omics and multiplexed imaging to explore cancer biology. Nature methods, 18(9):997–1012, 2021.

Mathieson, L. and Moscato, P. An introduction to proximity graphs. Business and Consumer Analytics: New Ideas, pages 213–233, 2019.

Moses, L. and Pachter, L. Museum of spatial transcriptomics. Nature methods, 19(5):534–546, 2022.

Qian, N. and Weinstein, J. A. Volumetric imaging of an intact organism by a distributed molecular network. bioRxiv, 2023.

Rodriques, S. G., Stickels, R. R., Goeva, A., Martin, C. A., Murray, E., Vanderburg, C. R., Welch, J., Chen, L. M., Chen, F., and Macosko, E. Z. Slide-seq: A scalable technology for measuring genome-wide expression at high spatial resolution. Science, 363(6434): 1463–1467, 2019.

Satija, R., Farrell, J. A., Gennert, D., Schier, A. F., and Regev, A. Spatial reconstruction of single-cell gene expression data. Nature biotechnology, 33(5):495–502, 2015.

Schaus, T. E., Woo, S., Xuan, F., Chen, X., and Yin, P. A dna nanoscope via auto-cycling proximity recording. Nature communications, 8(1):1–9, 2017.

Shanker, O. Defining dimension of a complex network. Modern Physics Letters B, 21(06): 321–326, 2007.

Shapiro, E., Biezuner, T., and Linnarsson, S. Single-cell sequencing-based technologies will revolutionize whole-organism science. Nature Reviews Genetics, 14(9):618–630, 2013.

Solomon, H. Geometric probability. SIAM, 1978.

Ståhl, P. L., Salmén, F., Vickovic, S., Lundmark, A., Navarro, J. F., Magnusson, J., Giacomello, S., Asp, M., Westholm, J. O., Huss, M., et al. Visualization and analysis of gene expression in tissue sections by spatial transcriptomics. Science, 353(6294):78–82, 2016.

Wang, G., Moffitt, J. R., and Zhuang, X. Multiplexed imaging of high-density libraries of rnas with merfish and expansion microscopy. Scientific reports, 8(1):1–13, 2018.

Watts, D. J. and Strogatz, S. H. Collective dynamics of ‘small-world’networks. nature, 393 (6684):440–442, 1998.

Weinstein, J. A., Regev, A., and Zhang, F. Dna microscopy: optics-free spatio-genetic imaging by a stand-alone chemical reaction. Cell, 178(1):229–241, 2019.

Yanai, I., Benjamin, H., Shmoish, M., Chalifa-Caspi, V., Shklar, M., Ophir, R., Bar-Even, A., Horn-Saban, S., Safran, M., Domany, E., et al. Genome-wide midrange transcription profiles reveal expression level relationships in human tissue specification. Bioinformatics, 21(5):650–659, 2005.

Zemel, R. and Carreira-Perpiñán, M. Proximity graphs for clustering and manifold learning. Advances in neural information processing systems, 17, 2004.

